# A Machine Learning-based approach for Simultaneous Detection of Interfering Analytes in Electrochemical Nanobiosensors

**DOI:** 10.1101/2024.03.11.584459

**Authors:** Ritwik Jain, Srishti Verma, Gorachand Dutta

## Abstract

Electrochemical biosensors can be used to detect analytes of importance precisely. These sensors generate rapid and accurate electrical signals that reveal the presence and concentration of the targeted analyte. Detecting multiple analytes simultaneously with an electrochemical biosensor is advantageous. It provides cost and time efficiency, multiplexing capability, and flexibility, making it valuable in diverse applications such as medical diagnostics, environmental monitoring, and industrial processes. However, simultaneous detection of analytes may suffer from the problem of interference. The interference effect causes the signal of an analyte at a particular concentration to deviate from the expected one. We observe a similar effect in the simultaneous detection of Folic Acid and Uric Acid using a nanomaterial-based electrochemical sensor. To address this effect, we propose a machine learning (ML) approach. ML algorithms handle complex interactions by autonomously identifying patterns, dependencies, and nonlinear relationships within data, enabling it to make predictions and decisions in intricate and dynamic scenarios. Our approach can be generalised to any two analytes showing interference and would scale well to interference between multiple analytes. We test several regression algorithms and compare their performance to the standard calibration plot method. As compared to the standard method, our approach shows a 4.49 µM decrease in concentration prediction error.

## I. Introduction

Electrochemical biosensors selectively react with a target analyte and produce an electrical signal related to the concentration of the analyte being studied [1]. As compared to traditional methods such as high performance liquid chromatography (HPLC), spectrometry, and fluorometry, these sensors are cheap, simple, and time-effective [2]. Electrochemical biosensors are often fabricated using nanomaterials that enhance the detection capabilities of the sensors [3], such sesnors are termed as nanobiosensors.

Some nanobiosensors can detect several important analytes at once [4], [5]. Such simultaneous detection capabilities are significant in clinical diagnostic settings where as much information as possible needs to be extracted from a single sample, saving time and cost, and allowing for a more robust diagnosis [6]. However, the simultaneous detection of two or more analytes in electrochemical sensors might suffer from the problem of interference [7], [8], [9]. This interference occurs due to the complex interactions between the analytes and the modified electrode surface. These interactions cause deviations in signal readouts so they can not be mapped to concentrations using linear methods. We observe an interference effect in the simultaneous detection of Folic Acid (FA) and Uric Acid (UA) using our nanobiosensor.

FA and UA are important molecules co-existing in some biological fluids [10]. A disruption in their concentrations could indicate diseases [11], [12]. Thus fast and reliable simultaneous determination of UA and FA is necessary.

Our work proposes a possible solution to resolve interference in electrochemical sensors using Machine Learning (ML) algorithms. The advantage of these algorithms is that they provide the opportunity for scaling up to the simultaneous detection of multiple interfering analytes by incorporating more input features. The pipeline is illustrated in Fig.1.

**Fig. 1:**
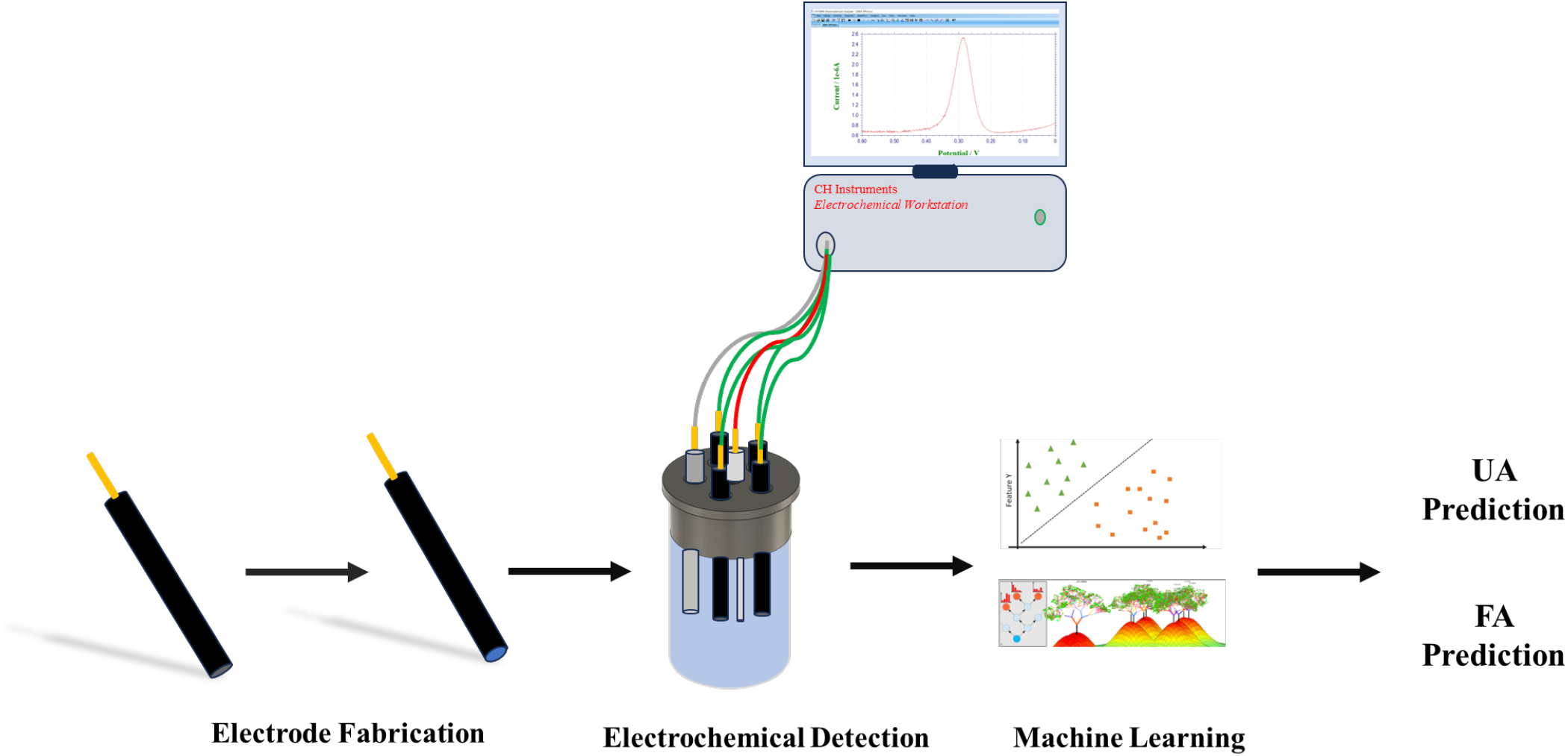
Pipeline for the machine learning approach

ML has been previously used in electrochemical biosensors, although its use has still been limited [13]. Aiassa et al. [14] and Rivera et al. [15] used ML to detect a single analyte to reduce the effect of background noise. Qian et al. [16], used statistical machine learning to quantify E.Coli concentration from water samples. In the case of interfering analytes, Shukla et al. [17] and Zhou et al. [18] used an artificial neural network (ANN) to quantify the concentration of target analytes. ANNs require large amounts of training data and moreover suffer from the blackbox problem, making them hard to interpret and be used in clinical settings [19]. We use the classical feature extraction followed by regression strategy to tackle these problems.

## II. Methods

### A. Electrode Fabrication

The fabrication process involved the modification of a glassy carbon electrode (3 mm diameter) surface. This modification ensures that the surface gets functionalised and in turn, enhances the detection of the analytes. This process involves mechanical and electrochemical cleaning followed by dropcasting the nanomaterial on the electrode surface. We used a novel nanomaterial which we will reveal and characterise in a future publication. Nanomaterials amplify analyte detection by providing a high surface area for increased interactions, enhancing sensitivity in various sensing technologies. Their unique properties lead to improved accuracy and efficiency in detecting trace amounts of analytes.

### B. Setup and Data Recording

A three-electrode system was used for this study. We used the modified electrode as the working electrode. The reference electrode was Ag/AgCl and the counter electrode was the Platinum wire. UA and FA were dissolved in 0.01 M Phosphate-buffered Saline (PBS), and the concentrations were determined.

We used the Differential Pulse Voltammetry (DPV) technique. DPV involves the measurement of current as a function of applied potential. Compared to cyclic voltammetry, DPV provides the advantages of having a higher signal-to-noise ratio and reducing the contribution of non-faradaic currents which, can interfere with the analysis. The readout of DPV and other voltammetry techniques is a current signal as a function of applied potential. The DPV data was recorded using a CHI Electrochemical Analyzer. An example of an output from DPV is shown in Fig. 2.

**Fig. 2:**
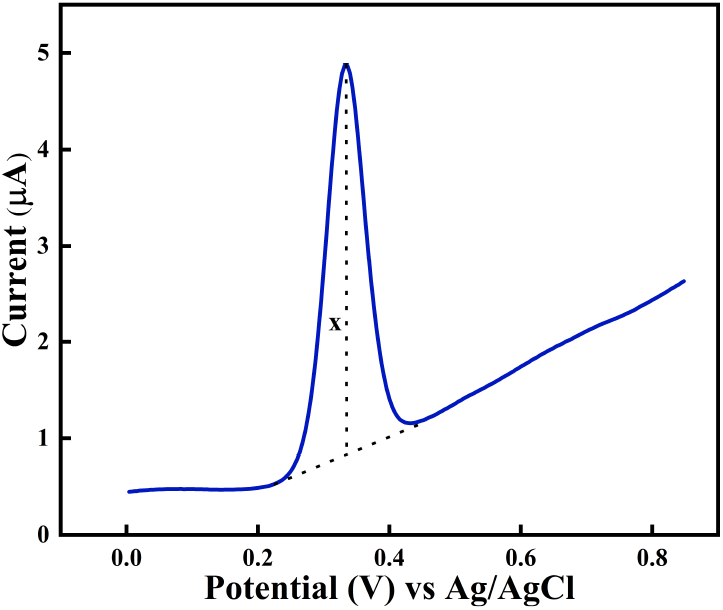
An example of output from a DPV. The peak, in this case corresponds to the oxidation of UA into Uric Allantoin. Here, ‘x’ signifies peak current.

### C. Machine Learning Algorithms and Input Features

We used a subclass of ML algorithms known as regression algorithms to predict the concentration. These algorithms aim to model the relationship between a dependent variable, which in this case would be the concentration, and one or more independent variables (features) by learning from training data [20]. These algorithms use an objective or loss function, quantifying the error between actual and predicted values. The goal is to minimize this function during training by optimising the parameters involved in the objective function. Different algorithms use different optimisation techniques, it is difficult to predict which algorithm would perform best for a particular dataset [21]. The performance of the regression algorithm is assessed using evaluation metrics such as Root mean squared error (RMSE), and Mean absolute error (MAE), which measure the accuracy and goodness of fit of the model to the data.

We used 4 popular regression algorithms: Random forest (RF), support Vector Machine (SVM), and K-Nearest neighbours (KNN). The hyperparameters for the algorithms were selected by performing a grid seach across an arbitrarily defined search space. The input features used were the peak heights corresponding to FA and UA, the baseline for calculating the peak heights was taken as shown in Fig.2. Our dataset comprised a total of 90 data points (DPV curves). These points range from UA concentrations 1 - 20 µM and FA concentrations of 5 - 50 µM. A high concentration (more than 5 µM) of FA was used because we had experimentally observed that the interference effect was negligible below this concentration. We used a five-fold cross-validation method to obtain the final prediction error.

## III. Results AND Discussion

### A. Interference effect

It is evident from Fig.3 that UA electro-oxidation peak occurs at 0.340 V, while in the presence of FA this peak shifts by 0.024 V towards higher potential. Electro-oxidation of FA occurs at 0.708 V. There is a significant gap in the peak potential of both the analytes but the addition of FA has an antagonistic effect on the detection of UA which means that the peak current of UA decreases upon addition of FA, but the peak current of FA remains constant with varying concentration of UA as shown in Fig.3.

**Fig. 3:**
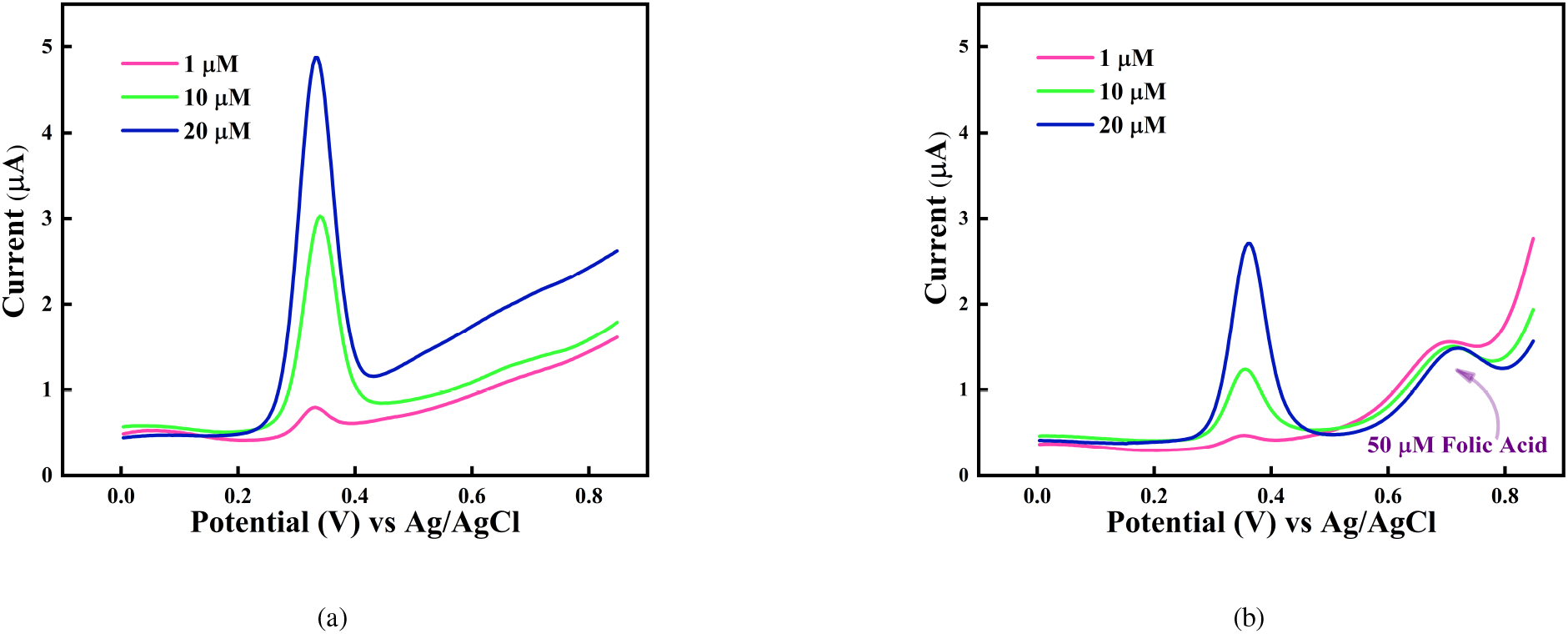
Diffrential pulse voltammetry responses for a) Uric acid at 1 - 20 µM b) Uric acid at 1 - 20 µM in the presence of 50 µM of Folic acid.

We have also observed that the decrease in the peak depends upon the concentration of FA added as evident in Fig.4(a) and 4(b). Similarly, it is observed that the peak of FA remains constant with varying UA concentrations (Fig. 4(c)).

**Fig. 4:**
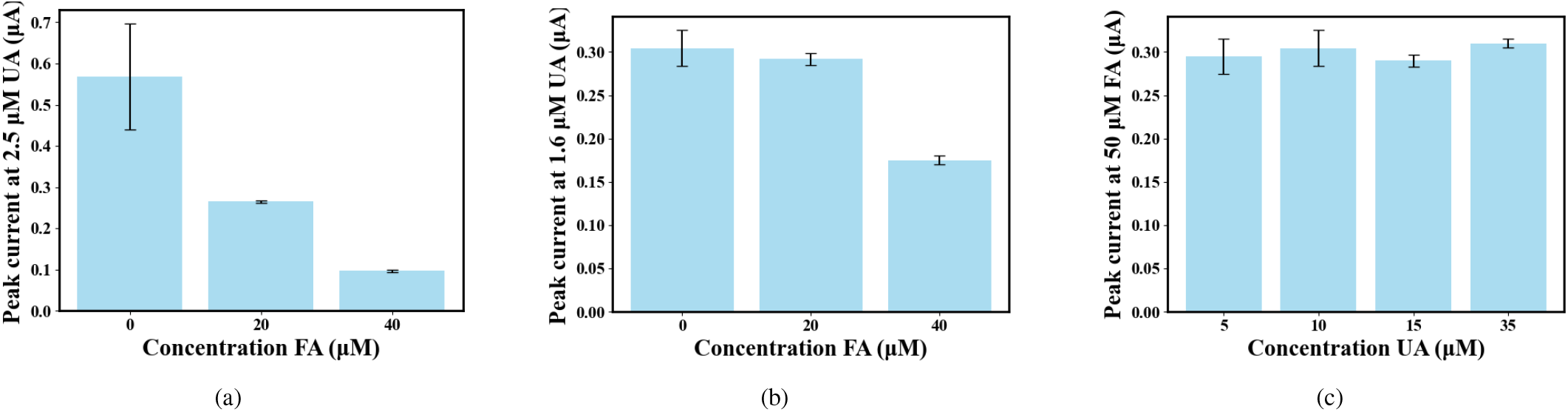
Peak Current dependence of FA and UA on each other. a) Peak current at 2.5 µM Uric acid when Folic acid concentration is varied b) Peak current 1.6 µM Uric acid when Folic acid concentration is varied c) Peak current at 50 µM Folic acid when Uric acid concentration is varied.

### B. Calibration Plots

The calibration plot for FA and UA have been shown in Fig. 5(a) and Fig. 5(b). When only either of the two analytes are present, their concentration can be found out using these calibration plots. FA concentration can be determined from this plot even in the presence of UA. The concentrations provided from these plots provides a baseline to which the performance of ML algorithms can be compared.

**Fig. 5:**
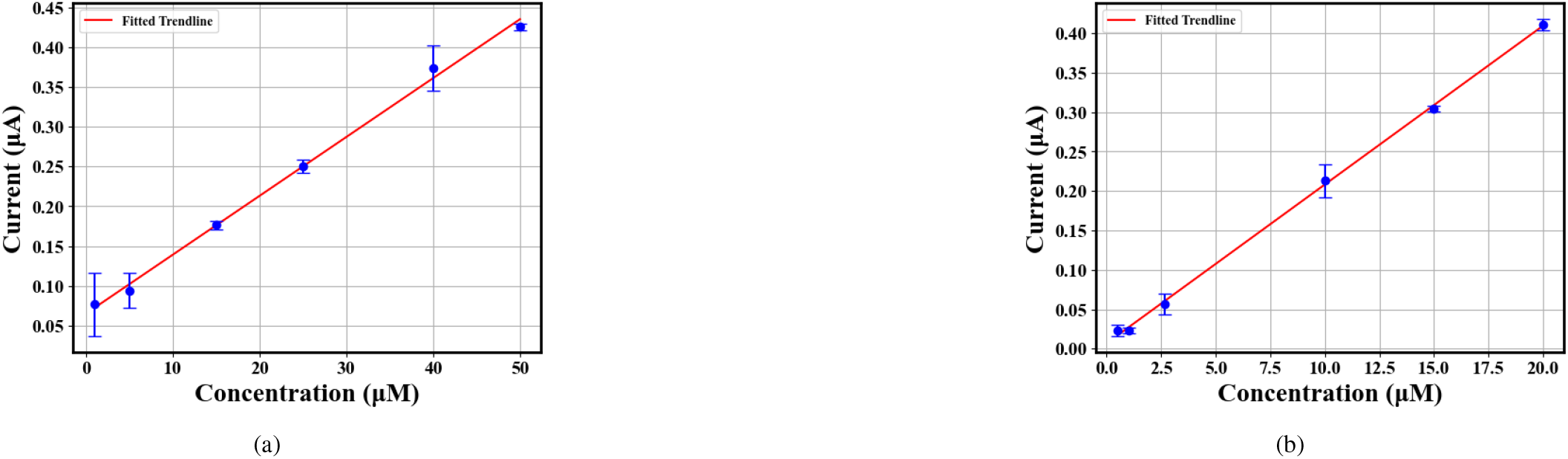
Calibration plots. a) Folic acid b) Uric acid.

### C. Performance of the Regression Algorithms

The RMSE and MAE of the dataset, as calculated from the calibration plot was 5.04 µM and 4.01 µM respectively. The lowest RMSE and MAE was obtained using the KNN regressor which had a decrease of RMSE and MAE of 4.49 µM and 3.59 µM respectively as compared to the calibration plot (Table I).

**TABLE 1:** Performance of different regression algorithms.

| Algorithm | RMSE ($\mu\text{M}$) | MAE ($\mu\text{M}$) |
| --- | --- | --- |
| SVM | 0.92 | 0.65 |
| Random Forest | 0.81 | 0.63 |
| KNN | 0.55 | 0.42 |
| XG Boost | 0.85 | 0.69 |

Fig.6 shows the RMSE for the KNN regressor across the five folds of four different UA concentration bounds in the dataset: 1-5 µM, 5-10 µM, 10-15 µM, 15-20 µM. The quality of the predictions was maintained throughout the range of UA concentrations in the dataset and the most accurate predictions were obtained when the UA concentration was between 5-10 µM.

**Fig. 6:**
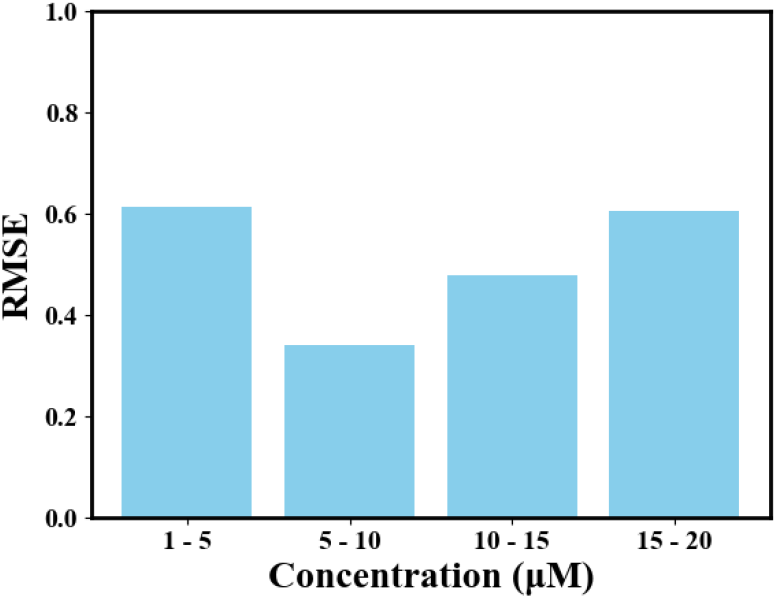
Prediction error for the KNN regressor corresponding to four different UA concentration bounds.

In our case of the simultaneous detection of two analytes we used only two input features (peak heights corresponding to UA and FA). However, in cases where there may be multiple analytes which are detected by the biosensor, multiple peak currents can be used as input features, although this would significantly increase the training data required by the ML algorithm. If the peaks of multiple analytes are merged then the peak potential or area under the curve can be taken as input features.

## IV. Conclusion AND Future Scope

In this study, we have proposed a generalisable machine learning-based approach to rectify the interference effect observed for the simultaneous detection of the analyte of importance. Here we have observed the interference of a higher concentration of Folic Acid on Uric Acid’s electrochemical detection. We used several regression algortihms to quantify the Uric acid concentration. K-Nearest neighbour regressor gave the best performance with an RMSE of 0.55 µM and showed a decrease in RMSE of 4.49 µM as compared to the calibration plot. A similar machine learning-based approach could be used to address the interference effect observed between multiple analytes, which could be the focus of future studies.

## References

[1] Niina J Ronkainen, H Brian Halsall, and William R Heineman. Electrochemical biosensors. Chemical Society Reviews, 39(5):1747–1763, 2010.

[2] Wei Zhang, Ruiguo Wang, Fang Luo, Peilong Wang, and Zhenyu Lin. Miniaturized electrochemical sensors and their point-of-care applications. Chinese Chemical Letters, 31(3):589–600, 2020.

[3] Naumih M Noah, Peter M Ndangili, et al. Current trends of nanobiosensors for point-of-care diagnostics. Journal of Analytical Methods in Chemistry, 2019, 2019.

[4] S Irem Kaya, Sevinc Kurbanoglu, and Sibel A Ozkan. Nanomaterials-based nanosensors for the simultaneous electrochemical determination of biologically important compounds: ascorbic acid, uric acid, and dopamine. Critical Reviews in Analytical Chemistry, 49(2):101–125, 2019.

[5] Aysun Savk, Buse Özdil, Buse Demirkan, Mehmet Salih Nas, Mehmet Harbi Calimli, Mehmet Hakki Alma, Abdullah M Asiri, Fatih Şen, et al. Multiwalled carbon nanotube-based nanosensor for ultrasensitive detection of uric acid, dopamine, and ascorbic acid. Materials Science and Engineering: C, 99:248–254, 2019.

[6] Greta Jarockyte, Vitalijus Karabanovas, Ricardas Rotomskis, and Ali Mobasheri. Multiplexed nanobiosensors: current trends in early diagnostics. Sensors, 20(23):6890, 2020.

[7] Wen-Zhi Jia, Kang Wang, and Xing-Hua Xia. Elimination of electrochemical interferences in glucose biosensors. TrAC Trends in Analytical Chemistry, 29(4):306– 318, 2010.

[8] Ananda Basu, Sona Veettil, Roy Dyer, Thomas Peyser, and Rita Basu. Direct evidence of acetaminophen interference with subcutaneous glucose sensing in humans: a pilot study. Diabetes technology & therapeutics, 18(S2): S2–43, 2016.

[9] Toshio Yao, Yoichi Taniguchi, Tamotsu Wasa, and Soichiro Musha. Anodic voltammetry and its analytical application to the detection and simultaneous determination of hypoxanthine, xanthine, and uric acid. Bulletin of the Chemical Society of Japan, 51(10):2937–2941, 1978.

[10] Tao Nie, Limin Lu, Ling Bai, Jingkun Xu, Kaixin Zhang, Ou Zhang, Yangping Wen, and Liping Wu. Simultaneous determination of folic acid and uric acid under coexistence of l-ascorbic acid using a modified electrode based on poly (3, 4-ethylenedioxythiophene) and functionalized single-walled carbon nanotubes composite. Int J Electrochem Sci, 8:7016–7029, 2013.

[11] F Jossa, E Farinaro, Salvatore Panico, V Krogh, E Celentano, R Galasso, M Mancini, and M Trevisan. Serum uric acid and hypertension: the olivetti heart study. Journal of human hypertension, 8(9):677–681, 1994.

[12] Alessio Di Tinno, Rocco Cancelliere, and Laura Micheli. Determination of folic acid using biosensors—a short review of recent progress. Sensors, 21(10):3360, 2021.

[13] Lucas B Ayres, Federico JV Gomez, Jeb R Linton, Maria F Silva, and Carlos D Garcia. Taking the leap between analytical chemistry and artificial intelligence: A tutorial review. Analytica Chimica Acta, 1161:338403, 2021.

[14] Simone Aiassa, Ivan Ny Hanitra, Gabriele Sandri, Tiberiu Totu, Francesco Grassi, Francesca Criscuolo, Giovanni De Micheli, Sandro Carrara, and Danilo Demarchi. Continuous monitoring of propofol in human serum with fouling compensation by support vector classifier. Biosensors and Bioelectronics, 171:112666, 2021.

[15] Elmer Ccopa Rivera, Jonathan J Swerdlow, Rodney L Summerscales, Padma P Tadi Uppala, Rubens Maciel Filho, Mabio RC Neto, and Hyun J Kwon. Data-driven modeling of smartphone-based electrochemiluminescence sensor data using artificial intelligence. Sensors (Basel, Switzerland), 20(3), 2020.

[16] Hanyu Qian, Eric McLamore, and Nikolay Bliznyuk. Machine learning for improved detection of pathogenic e. coli in hydroponic irrigation water using impedimetric aptasensors: a comparative study. ACS omega, 8(37): 34171–34179, 2023.

[17] Rajendra P Shukla, Matan Aroosh, Alon Matzafi, and Hadar Ben-Yoav. Partially functional electrode modifications for rapid detection of dopamine in urine. Advanced Functional Materials, 31(17):2004146, 2021.

[18] Zhongzeng Zhou, Luojun Wang, Jing Wang, Conghui Liu, Tailin Xu, and Xueji Zhang. Machine learning with neural networks to enhance selectivity of nonenzymatic electrochemical biosensors in multianalyte mixtures. ACS Applied Materials & Interfaces, 14(47):52684–52690, 2022.

[19] Zhongheng Zhang, Marcus W Beck, David A Winkler, Bin Huang, Wilbert Sibanda, Hemant Goyal, et al. Opening the black box of neural networks: methods for interpreting neural network models in clinical applications. Annals of translational medicine, 6(11), 2018.

[20] Rishabh Choudhary and Hemant Kumar Gianey. Comprehensive review on supervised machine learning algorithms. In 2017 International Conference on Machine Learning and Data Science (MLDS), pages 37–43. IEEE, 2017.

[21] Ritwik Jain, Prakhar Jaiman, and Veeky Baths. Feature engineering for an efficient motor related ecog bci system. In 2023 IEEE International Conference on Systems, Man, and Cybernetics (SMC), pages 4720–4727. IEEE, 2023.

